# High yield purification of an Isoleucine zipper modified CD95 Ligand with either biotin or DNA-oligomer binding domain for efficient Cell Apoptosis Induction

**DOI:** 10.1101/2024.10.12.618001

**Authors:** Xiaoyue Shang, Nina Bartels, Johann Moritz Weck, Sabine Suppmann, Jérôme Basquin, Amelie Heuer-Jungemann, Cornelia Monzel

## Abstract

**Background:** Cluster of differentiation 95 (CD95/Fas/Apo1) as part of the Tumor-necrosis factor (TNF) receptor family is a prototypic trigger of the ‘extrinsic’ apoptotic pathway and its activation by the trimeric ligand CD95L is of high interest for anticancer therapy. However, CD95L, when presented in solution, exhibits a low efficiency to induce apoptosis signaling in human cells.

**Results:** Here, we design a recombinant CD95L exhibiting an isoleucine zipper (IZ) motif at the N-terminus for stabilization of the trimerized CD95L and demonstrate its high apoptosis induction efficiency. A cysteine amino acid fused behind the IZ is further used as a versatile coupling site for bionanotechnological applications or for the development of biomedical assays. A fast, cheap, and high-yield production of CD95L *via* the HEK293T secretory expression system is presented, along with CD95L affinity purification and functionalization. We verified the biological activity of the purified protein and identified a stabilized trimeric CD95L structure as the most potent inducer of apoptosis signaling.

**Conclusions:** The workflow and the findings reported here will streamline a wide array of future low- or high-throughput TNF-ligand screens, and their modification towards improving apoptosis induction efficiency and anticancer therapy.

## Background

CD95 ligand (CD95L, Fas ligand, CD178, or TNFSF6) is a 40 kDa type II transmembrane protein belonging to the tumor necrosis factor (TNF) superfamily (1). It is expressed on multiple immune cells such as cytotoxic T lymphocytes and monocytes (2). CD95L is the only ligand binding to the receptor CD95 (Fas, Apo-1), which upon ligand activation triggers the programmed cell death of apoptosis (3). Soon after the discovery of this apoptosis-induction mechanism, the CD95L/CD95 interaction emerged as a potential anticancer therapy (4). Yet accumulating evidence indicates another role of CD95L/CD95 in alternative nonapoptotic signaling pathways leading to proliferation or migration, and consequently, tumorigenesis (5,6). While activation of the proliferation pathway was suggested to involve additional interactions on the intracellular site, such as tyrosine phosphorylation of the intracellular CD95 death domain or of caspase-8, both by src-family kinases (SFKs) (7–10), the efficiency of apoptosis induction appears to also largely depend on the presentation of the CD95 ligand: for example, CD95L can exist in a membrane-anchored (mCD95L) as well as in a soluble form (sCD95L). In case of the soluble form, CD95L is cleaved by matrix metalloproteases (MMPs) at its stalk region (11) (see **Figure 1A**). Several studies showed that this soluble form of CD95L is highly inefficient in inducing apoptosis in comparison to mCD95L and manifested the idea that the CD95L molecular structure plays a decisive role (1): some studies reported that the metalloprotease-cleaved CD95L engenders a homotrimer, which is unable to induce cell death in certain pathologies, but instead triggers pro-inflammatory signaling pathways (12–16). Other studies showed, that soluble CD95L gets accumulated and, once it reaches a sufficient aggregation level with bound CD95, allows the implementation of the cell death program (17,18). Similarly, a recombinant form of two trimers was efficiently inducing cell death (19), supporting that the extent to which CD95L is multimerized is a pivotal step in determining whether cell apoptosis signaling is induced.

**Figure 1.**
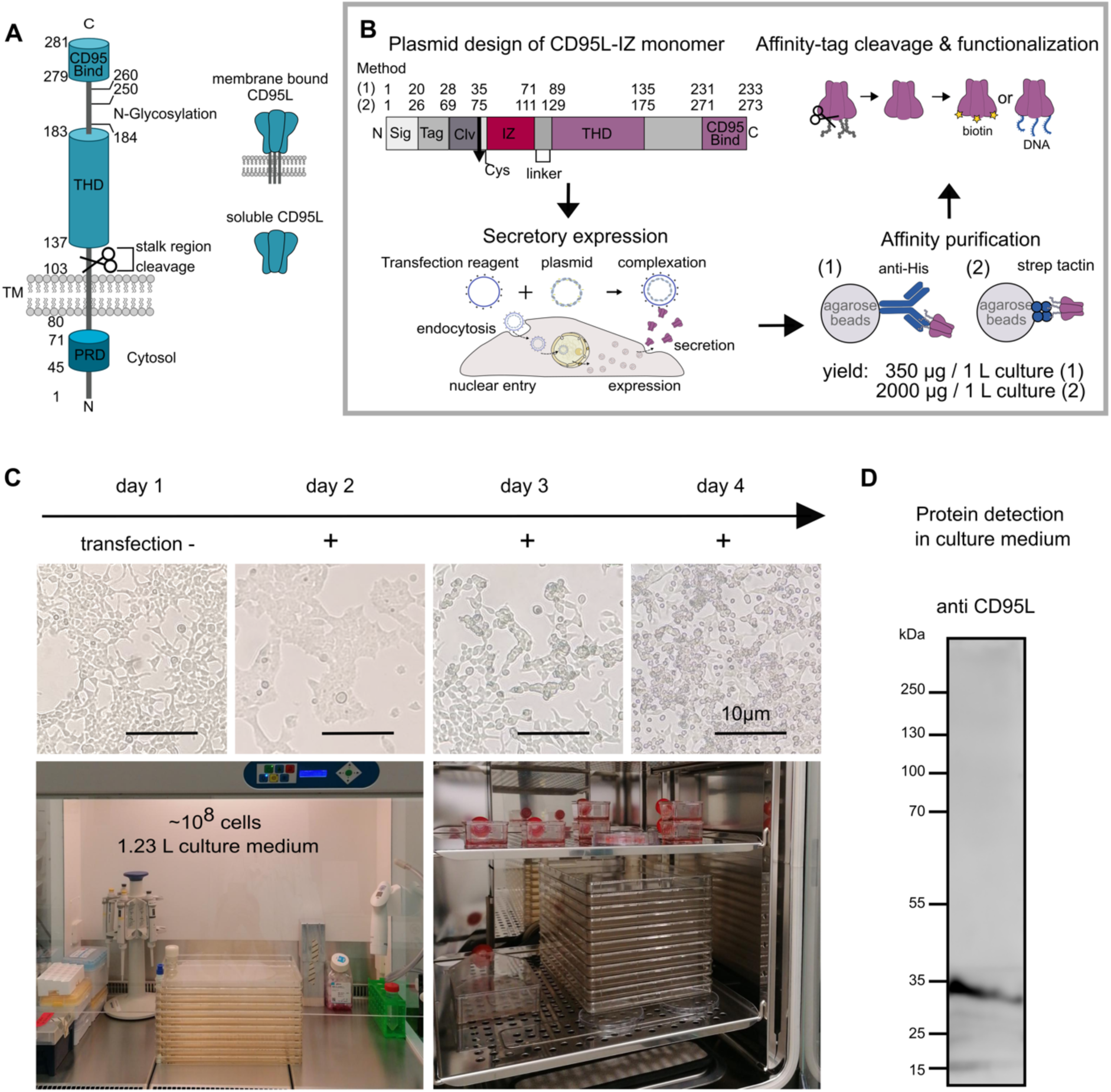
CD95L structure and experimental design. **A** CD95L can exist either in a membrane-bound or in a soluble form, the latter resulting from protease cleavage. Key domains include the CD95 binding domain (CD95 Bind), the TNF homology domain (THD) for its homotrimerization, and the proline-rich domain (PRD). **B** Workflow of the CD95L-IZ construct design, secretory expression, affinity purification, and modification. The recombinant protein consists of a signaling peptide (Sig), an affinity tag (Tag), a cleavage site (Clv), a Cysteine (Cys), an isoleucine zipper domain (IZ) fused via a linker to the extracellular CD95L domain. In method 1 transient CD95-IZ expression was used. Here, a His-tag was used as Tag and a TEV protease sequence as Clv. Affinity purification was done by anti-His-tag antibody coupled agarose beads. In method 2 a cell line stably expressing CD95-IZ was used. Here, a Twin-Strep-Tag was used as Tag and a PreScission cleavage site as Clv. Affinity purification was done by Strep-TactinXT4Flow beads. **C** Upper panel: examination of cell morphology before and up to three days after transfection. Lower panel: photos of cell culture and transfection using a 10-layer cell factory in a cell culture hood (left) or incubator (right). A T175 cell culture flask was seeded with the same cell density as in a 10-layer cell factory to check the cell condition and confluency. **D** CD95L-IZ protein expression and secretion was verified by a Western blot of the cell culture medium using anti-CD95L antibody. The image contrast was adjusted for better visibility.

CD95L has a long cytoplasmic domain with its N-terminus, a transmembrane (TM) domain, a stalk region, a TNF homology domain (THD) that mediates homotrimerization (1), and a CD95 binding domain at its C-terminus (see **Figure 1 A**). Both, the membrane-anchored and soluble forms of CD95L, form homotrimers (12). However, the studies mentioned above suggest that the binding of naturally occuring sCD95L to CD95 exhibits a reduced efficiency to induce cell death (14), due to a missing secondary stabilization of the trimer at the stalk region compared to the membrane-bound CD95L. Indeed, several structural modifications or intermolecular crosslinking approaches have been tested to probe and improve the sCD95L apoptosis efficiency. For example, the fusion of an Fc domain (19) or the crosslinking of sCD95L with an antibody (20) showed, that a significant enhancement leading to some 38-81 % of apoptosis in single cells *in vitro* in comparison to 15-26 % of the soluble form can be obtained. Since CD95L is often discussed as a target in anti-cancer treatment, increasing the CD95L efficacy is of high interest in anti-cancer drug development.

In this work, we use a variant of the leucine zipper (LZ) as a self-crosslinking domain of CD95L, as it is a dimerization motif/coiled-coil motif found in eukaryotic transcription factors (21). Each monomer forms an alpha helix with a periodic repetition of leucine residues at every seventh position. These motifs are characterized by a heptad repeat containing hydrophobic residues at the first (a) and fourth (d) positions, facilitating hydrophobic interactions and dimerization (22). The variant of the wild-type GCN4 leucine zipper used, is the isoleucine zipper (IZ), which adopts parallel trimer structures with isoleucine substitutions at all a and d positions (23,24). The isoleucine zipper is a relatively small and straightforward motif, which can be advantageous when minimizing the overall size of the fusion protein. This facilitates easier expression, folding, and purification of the protein compared to larger domains like the T4 foldon or tenascin-C. IZ trimerization has been used to enhance the biological activity of CD40L (25,26) or TNFR receptor proteins (27,28). Our goal in this study is to introduce the IZ domain to CD95L to enhance apoptosis efficiency without additional crosslinkers and to obtain new insights into the mechanism of apoptosis signal induction.

To tailor CD95L constructs towards their efficient apoptosis induction requires their genetic modification and the appropriate expression system to obtain the protein cost-effectively and at a high yield. Soluble CD95L expression and purification have been widely studied in both eukaryotic (*Dictyostelium discoideum*, COS cells) (29–32) and prokaryotic expression systems, such as *Escherichia coli* (*E.coli*) (33). The bacterial *E.coli* expression system has been the most popular method for recombinant protein expression and purification, due to the benefits of cost-efficiency, time-saving, and high yield at the milligram scale (34). However, post-translational modifications such as disulfide-bond formation and glycosylation are very different in prokaryotes compared to mammalian cells (35).Previous studies have demonstrated that mutations at CD95L glycosylation sites do not significantly affect self-aggregation or receptor binding (36). However, these mutations reduce the CD95L expression level in transfectants (37), indicating a positive relation between glycosylation and protein maturation and stability. Glycans can also protect proteins from proteolytic degradation by masking protease sites, thereby prolonging the half-life of glycoprotein at the cell surface (38).

While low yield has always been the cutting limit of human protein expression and purification using an adherent cell line, the most appropriate host to maintain the native folding environment and modifications is the human cell line HEK293 (human embryonic kidney 293). In the past, also the adapted suspension cell line HEK293F as a fast-growing variant has been established (39). Another variant, which was used in this work, is the HEK293T cell line, a highly transfectable derivative of the HEK293 cell line, due to the incorporation of a temperature-sensitive mutant of SV40 large T-antigen (tsA1609). This cell line maintains a high copy number of transfected plasmid DNAs which carry the SV40 origin of replication (40).

Since CD95L in the form of an efficient apoptosis inducer is of high interest in anti-cancer drug development, we here designed a CD95L fused with an isoleucine zipper domain (CD95L-IZ), which enhances apoptosis efficiency without the need for additional crosslinking. We show that we can produce a large batch of CD95L-IZ using the mammalian HEK293T cell line as an expression system and the cell factory system for the cell culture, reducing the production time from months to one week. We first used a transient expression system followed by immunoprecipitation purification via an anti-His affinity resin (method 1). We then used the findings from method 1 to develop an alternative strategy based on a stable cell line expression and purification via a strep-tactin resin to increase the yield and to simplify the cell cultivation process (method 2). In both cases we obtained CD95L-IZ at high purity and yield (see **Methods** for details). We characterize the CD95L-IZ oligomeric state, and demonstrate its functionality to create bionanotechnological or biomedical assays with a high efficiency to induce apoptosis in cancer cells.

## Methods

Most of the presented data have been acquired using CD95L-IZ from transient transfections of HEK293T cells. In addition, a HEK293T cell line was created stably expressing CD95L-IZ. Here, we compare describe the handling in both cases along with the expression efficiencies. First the methods for transient transfections are described followed by methods for the stable cell line expression.

### CD95L-IZ plasmid design for transient expression

For the plasmid IL2pept-His8-TEV-Cys-IZ-CD95L (137–281), the extracellular domain of CD95L consisting of amino acids 137-281, was fused at its N-terminus with an IZ domain, which supports and stabilizes the trimerization of CD95L. One extra cysteine was added after the IZ for further functionalization (biotinylation or DNA conjugation). Next to the cysteine, a TEV protease cleavage site ENLYFQ was incorporated along with 8 histidines, to allow for His-tag purification. To allow protein secretion, we used the interleukin 2 (IL2) signal peptide MRRMQLLLLIALSLALVTNS as the N-terminal sequence, which has been extensively studied and widely used in human protein production in industry and academia (41–43). Specifically, we used a mutant version of the IL-2 signal peptide (IL2pept) with modifications in the basic and hydrophobic domains, which resulted in a three-fold increase in protein secretion levels compared to the wild type (44). Notably, secretory expression levels are often higher than those of intracellular soluble protein expression due to more efficient protein folding and processing, reduced proteolytic degradation and cellular stress, and enhanced protein stability. The signal peptide is a short amino acid sequence located at the N-terminal of the nascent protein and directs the secretion pathway. The complete amino acid sequence of the insert reads MRRMQLLLLIALSLALVTNSHHHHHHHHENLYFQGCGDRMKQIEDKIEEILSKIYHIENEIARIKKLIGERTSGGSGGTGGSGGTGGSPPEKKELRKVAHLTGKSNSRSMPLEWEDTYGIVLLSGVKYKKGGLVINETGLYFVYSKVYFRGQSCNNLPLSHKVYMRNSKYPQDLVMMEGKMMSYCTTGQMWARSSYLGAVFNLTSADHLYVNVSELSLVNFEESQTFFGLYKL*. It has a theoretical isoelectric point and molecular weight pI / M.W. of: 9.10 / 26312, respectively, as calculated by using Expasy (https://web.expasy.org/compute_pi/). The corresponding DNA was inserted into a plasmid with a pcDNA3.1(-) backbone with a 5‘ restriction site EcoRI and 3‘ restriction site BamHI. The plasmid was ordered from BioCat (BioCat GmbH, Heidelberg, Germany) with codon optimization for CD95L-IZ-His protein expression in human cell lines.

### CD95L-IZ plasmid amplification

For large-scale CD95L-IZ-His expression and purification using transient transfection, a milligram amount of plasmid DNA was required. The plasmid was purified using a NucleoBond® Xtra Maxiprep kit (Macherey-Nagel, Düren, Germany). The plasmid was transformed into DH5*⍺* competent cells (Thermo Fisher Scientific, Waltham, Massachusettes, USA) *via* heat shock and spread on LB (lysogeny broth) agar plates with ampicillin for resistant strain selection and incubated overnight at 37 °C. Thereafter, a single colony was picked and a preculture in 5 ml 2 x YT (yeast extract tryptone) medium supplemented with ampicillin for 5 h was started. The preculture was distributed over 4 x 500 ml of 2 x YT medium and *DH*5*⍺* cells were grown in a shaker at 37 °C and 200 rpm (INFORS HT, Bottmingen, Switzerland) overnight. From each 500 ml culture, 2 ml of purified plasmid at a concentration of 1.4-1.9 µg/µl was obtained. The correct plasmid replication was checked by DNA sequencing. Of note, for continuous CD95L-IZ production, the establishment of a CD95L-IZ stably expressing cell line is recommended, since no large amount of plasmid DNA is required and cells can be sorted for those exhibiting a high transfection efficiency. The formation of such stable cell line is described below in the subsection **formation of stable cell line and protein purification**.

### Cell culture

Hela wild type cells (Hela WT) and human embryonic kidney 293 T (HEK293T) cells were ordered (both from ATCC® CCL-2™, ATCC, Manassas, VA, USA). Hela CD95 knock out (Hela CD95^KO^) stable cell line was generated using clustered regularly interspaced short palindromic repeats (CRISPR/Cas9) (45), the guide RNA was CATCTGGACCCTCCTACCTC (46). All cell lines were cultured in Dulbecco’s modified eagle medium (DMEM) + GlutaMAX^TM^ (Gibco, Life Technologies Inc., Carlsbad, CA, USA) containing 10 % fetal bovine serum FBS (Gibco) and 1 % penicillin/streptomycin (P/S) solution (Sigma-Aldrich, Merck KGaA, Darmstadt, Germany) at 5 % (v/v) CO_2_ in a humidified incubator (PHCbi, PHC Holdings Corporation, Tokyo, Japan). Cells were maintained in T25 flasks (Sarstedt AG & Co. KG, Nümbrecht, Germany) and the cell morphology and sterile conditions were checked regularly on the microscope. Cells were passaged every 2-3 days at a confluency level of around 60-80 %, as follows: i) cells were detached with trypsin-EDTA solution (Sigma) incubated for 4-5 min, ii) 5 ml of complete medium was added to inhibit the trypsin reaction, iii) cells were centrifuged at 1300 g for 5 min at room temperature (r.t.) to remove the trypsin containing supernatant, and iv) cells were resuspended in the fresh medium and seeded afterwards.

### Large-scale expression of CD95L-IZ using cell factory

The adherent HEK293T cell line was used to increase the recombinant protein production yield. Before large-scale cell culture and transfection, HEK293T cells were tested for mycoplasma contamination and, if necessary, treated with gentamicin (Sigma-Aldrich) at 10 µg/ml for a week to eradicate mycoplasma. A small-scale test expression was performed in a T75 flask and the protein expression and secretion were verified before starting the large-scale protein production. The large-scale cell culture and expression of CD95L-IZ-His was performed using a 10-layer cell culture vessel (Nunc™ EasyFill™ Cell Factory™ Systems, Thermo Fisher Scientific). Before using the cell factory, the number of cells needed to be scaled up to ∼ 10^!^ to start the culture at sufficient cell density. To this end, cells were first grown in a T25 flask, then transferred into a T175 flask and grown nearly to confluency. After 3 days, the up to 2.3 × 10^“^ cells which can be grown in a T175 flask, were distributed among seven T175 flasks and again grown to near confluency (80-90 %). Before the transfer, 1.23 L of DMEM complete (10 % FBS +1 % P/S) was filled into the 10-layer cell factory under sterile conditions and all cells were evenly distributed among the layers. Since it is difficult to monitor the cell growth in the 10-layer cell factory, another T175 flask was prepared in parallel at the same cell surface coverage. After 3 days of growth, when cells reached a confluency of 70-80 %, the DMEM complete medium in the cell factory was carefully replaced with the Opti-MEM complete (10 % FBS +1 % P/S) medium. Care should be taken to keep the inlet of the flasks dry to reduce the risk of contamination. For transfection, 1.28 mg of plasmid DNA was diluted into 60 ml Opti-MEM plain, and 3.84 mg of PEI was diluted into 60 ml Opti-MEM plain (DNA/PEI mass ratio of 1:3), the solutions were mixed and the 120 ml transfection mixture incubated at r.t. for 30 min. Afterwards, the mixture was added into a 1.23 L fresh Opti-MEM complete medium. Cell morphology and sterile conditions were checked every day after transfection. 72 h after the protein expression started, 1.35 L of cell culture medium was harvested. On average, this yielded an amount of 350 µg purified protein per 1 L culture medium.

### His-tag affinity purification

After cell culture medium collection, the cell debris was removed by centrifugation at 1300 g for 5 min. Large particles were removed by filtration with a 0.2 µm cut-off filter (Filtropur S 0.2, Sarstedt, Nümbrecht, Germany). The cell culture medium was concentrated using Amicon® Ultra-15 3 K centrifugal filters (Merck Millipore Ltd., Burlington, MA, USA) into a final volume of 80 ml. Thus, the ligand concentration in the culture medium was increased and the binding efficiency to the anti-His-tag affinity resin in the following step improved. Purifying His-tagged proteins using Ni-NTA techniques can be challenging, especially when the target protein is a small fraction of the sample. As an alternative, anti-His affinity resin has proven effective in antibody and mammalian protein purification at low expression levels. Therefore, we used agarose beads coupled with anti-His-tag antibody for CD95L-IZ-His purification. For purification of CD95L-IZ-His, 2 ml of anti-His-tag affinity resin (GenScript Biotech, Piscataway, NJ, USA) was added into the concentrated culture medium and incubated at 4 °C overnight with rotation. Thereafter, two empty polypropylene 1 ml columns (QIAGEN GmbH, Hilden, Germany) were equilibrated with 5 ml of 20 % ethanol, 5 ml of ultrapure water, and 5 ml TBS buffer (Tris-buffered saline, 50 mM Tris·HCl, 150 mM NaCl, pH 7.4). After the overnight incubation, the resin-culture medium mixture was loaded on both columns (each column volume: 40 ml). The loaded column was washed with 30 ml TBS buffer to rigorously remove non-specifically bound protein. Considering the theoretical pI of CD95L of 8.8 (for the His-tagged construct but without the cleaved signaling peptide after secretion), the CD95L-IZ-His was eluted following the GenScript manual using alkaline elution buffer (0.1 M Tris·HCl, 0.5 M NaCl, pH 12.0) with 12 ml total elution volume. After additional concentrating the 12 ml elution using Amicon® Ultra-15 3 K, the CD95-IZ-His concentration was determined by Pierce^®^ BCA Protein Assay Kit (BCA assay, Thermo Fisher Scientific). This yielded a protein concentration of 0.409 mg/ml. The protein size and purity were checked by SDS-PAGE using 10 % precast gel (Mini-PROTEAN® TGX™, Bio-Rad, Hercules, CA, USA). Here, 1 µg purified protein was loaded into 25 µl Laemmli sample buffer containing 50 mM TCEP·HCl and heated at 95 °C for 5 min followed by 180 V electrophoresis for 1 h. The gel was stained with SimplyBlue^TM^ Safe Stain (Thermo Fisher Scientific) according to the manufacturer’s protocol.

### Size exclusion chromatography

Size exclusion chromatography was performed using the Superdex 75^TM^ 10/300 GL (Cytiva) column with the ÄKTA prime plus (Cytiva, Marlborough, MA, USA). The column and system were equilibrated with 2 column volumes (CV) of 20 % ethanol, 2 CV of water, and 2 CV of phosphate-buffered saline (PBS) buffer at pH 7.4. Either 350 µl of purified CD95L-IZ-His (0.409 mg/ml) or a standard protein mix at 5.5 mg/ml (Sigma-Aldrich) was loaded and eluted at 0.3 ml/min flow rate with 0.3 ml fraction size in PBS buffer. Collected protein fractions were analyzed by SDS-PAGE. The standard protein mix included the following proteins: thyroglobulin bovine (670 kDa), γ-globulins from bovine blood (150 kDa), ovalbumin (44.3 kDa), ribonuclease A type I-A (13.7 kDa), and p-aminobenzoic acid (pABA) (137 Da).

### Western blot

To verify CD95L-IZ-His was present in the cell culture medium after cell transfection, a western blot was performed. Because the expression level was estimated to be in the ng/ml to ug/ml range, the medium needed to be concentrated by a factor 20-30 using Amicon Ultra-15 3 K centrifuge filters (Sigma-Aldrich). 20 µl of the concentrated medium was reduced with 5 µl of 5 x sample buffer with TCEP·HCl without heating in order to prevent FBS precipitation, which would lead to a high sample viscosity. Proteins were first separated by vertical electrophoresis (Mini-PROTEAN Tetra Vertical Electrophoresis Cell, Bio-Rad) with tris-glycine-SDS running buffer (Thermo Fischer Scientific) at 130-200 V. Thereafter, for western blot, a ROTI®PVDF membrane (Carl-Roth GmbH & Co. KG, Karlsruhe, Germany) was activated with 100 % methanol for 1 min, distilled water for 2 min and blotting buffer (25 mM Tris, 192 mM Glycine, pH 8.3, 20 % methanol) for 5 min. The membrane was sandwiched by two pieces of thick filter paper (diameter 110 mm, prewet in blotting buffer for 5 min) and the protein was transferred using a semi-dry system (Trans-Blot Turbo Transfer System, Bio-Rad) at 25 V 2.5 A for 20 min. The membrane was blocked with 5 % albumin fraction V BSA (ITW Reagents Pancreac, Barcelona, Spain) in TBST buffer (tris-buffered saline, 0.1 % tween-20) at r.t. for 1 h and probed with primary antibody at 4 °C overnight with 1:500 anti-His-tag antibody (BioLegend, J099B12) and 1:200 anti-CD95L antibody (Abnova, 2G9-G8). The next day, after rigorous washing 4 × 10 min each with TBST, the membrane was treated with goat anti-mouse (1:1000) or goat anti-rabbit (1:1000) secondary antibody coupled with horseradish peroxidase (HRP) (Cell Signaling Technology, Danvers, MA, USA) at r.t. for 60 min. Chemiluminescence detection was achieved by treating with Westar eta c ultra 2.0 substrate (Cyanagen Srl, BO, Italy) for 10-30 s. Images were taken using the Amersham^TM^ Imager AI680 (Cytiva).

### His-tag cleavage

200 µg of freshly eluted CD95L-IZ-His was reduced with 5 mM dithiothreitol (DTT) and incubated with 13 µg (1:15 w/w) TEV protease (Sigma-Aldrich) at 4 °C overnight. The next day, 50 µl of HisPur™ Ni-NTA magnetic beads (Thermo Fisher Scientific) were added in a 1.5 ml Eppendorf tube and washed with first 200 µl and then 500 µl of PBS buffer by mixing followed by collecting the beads with a magnetic stand and discarding the supernatant. Thereafter, 200 µg of the protein mixture was added and mixed by pipetting for 10 s. For coupling the CD95L-IZ-His to the beads, the mixture was rotated for 30 min at r.t. The beads were magnetically trapped and the supernatant containing CD95L-IZ without any His-tag was collected and checked by western blot. 60 ng of CD95L-IZ-His and CD95L-IZ after His-tag cleavage was loaded first on a 10% SDS-PAGE followed by a western blot probed with anti-His-tag antibody 1:1000 (BioLegend, San Diego, CA, USA). After His-tag cleavage, DTT must be removed due to the competition of coupling sites with maleimide. This was achieved by buffer exchange with PBS buffer using Amicon® Ultra-0.5 3 K centrifugal filters (Merck Millipore Ltd.).

### Formation of stable cell line and protein purification

HEK293T (ATCC® CRL-3216™) cell pools stably expressing Twin-Strep-tagged CD95L-IZ fusion protein were generated using piggyBac transposase as described previously (47). This expression system uses three components: the tetracycline-inducible expression vector PB-T-PAF, the helper plasmid PB-RN for transactivation of the Tet promotor and the plasmid expressing hyperactive piggyBac transposase under the control of a CMV promotor. To ensure efficient secretion, the human VEGF leader sequence (MNFLLSWVHWSLALLLYLHHAKWSQA) was used followed by a N-terminal Twin-Strep-Tag in combination with a PreScission cleavage site LEVLFQ/GP for tag removal. The complete amino acid sequence of the insert reads MNFLLSWVHWSLALLLYLHHAKWSQAAPMAEGGGQNSAWSHPQFEKGGGSGGGSGGSAWSHPQFEKTAGLEVLFQGPGCGDRMKQIEDKIEEILSKIYHIENEIARIKKLIGERTSGGSGGTGGSGGTGGSPPEKKELRKVAHLTGKSNSRSMPLEWEDTYGIVLLSGVKYKKGGLVINETGLYFVYSKVYFRGQSCNNLPLSHKVYMRNSKYPQDLVMMEGKMMSYCTTGQMWARSSYLGAVFNLTSADHLYVNVSELSLVNFEESQTFFGLYKL*.

Briefly, PB-T-PAF-IZ-CD95L under the control of an inducible Tet-Promotor was co-transfected with PB-RN and pCMV-HypBase, selected with puromycin for three weeks, and further expanded without selection. For protein production, stably CD95L-IZ expressing HEK293T cell pools were grown to a density of 1 × 10^6^ cells / mL in HEK FreeStyle293 medium (Gibco, Thermo Fisher Scientific), gene expression was induced using 1 µg/mL doxycycline and culture supernatant was harvested three days post-transfection.

For CD95L-IZ purification (see **Supplementary Figure 7**), 2 L prefiltered culture supernatant was treated with 4 mL BioLock (IBA LifeSciences, Göttingen) for 15 min on ice to block medium biotin content and subsequently loaded on 2 mL Strep-TactinXT4Flow beads (IBA LifeSciences, Göttingen). Beads were washed with 20 mL PBS and protein was eluted in PBS + 50 mM biotin. Eluted fractions were pooled for overnight in-solution cleavage at 4°C with HRV-3C-Protease (His-3C; Sigma-Aldrich). His-3C was depleted on the His-Select Affinity matrix (Sigma-Aldrich) prior to Size Exclusion Chromatography on a HiLoad 16/60 Superdex200 column (Cytiva, Germany) in PBS. Major peak fractions were analyzed by SDS-PAGE (see **Supplementary Figure 7**), pooled and concentrated using Vivaspin Turbo at 10 kDa cutoff (Sartorius Stedim Biotech, Göttingen) to a final concentration of 0.4 mg/mL. The final yield was 4 mg / 2 L culture.

### Site-specific labelling (biotinylation or DNA conjugation)

100 µg of protein in PBS was reduced freshly with 2 mM TCEP·HCl at 4 °C for 30 min. For biotinylation, 10 x molar excess of biotin-C5-maleimide (Sigma-Aldrich) in comparison to the protein was incubated with CD95L-IZ at 4 °C o/n (i.e. 2 µl of 10 mg/ml biotin-C5-maleimide was added). The next day, extra biotin was removed using NAP™-5 column (Cytiva). Here, the NAP-5 column was washed with 10 ml PBS buffer and then loaded with 120 µl of protein/biotin. After equilibration of the column with 0.38 ml PBS, the protein was eluted with 0.53 ml PBS. The final concentration of the biotinylated protein was 0.189 mg/ml in 550 µl. Biotinylation was checked *via* western blot: 300 ng of CD95L-IZ and CD95L-IZ-biotin was analyzed on 10 % SDS-PAGE followed by western blot and probed first by AF488-streptavidin (AAT Bioquest, Pleasanton, CA, USA) binding at 30 µg/ml overnight at 4 °C. Equal loading of CD95L-IZ/CD95L-IZ-biotin was qualitatively checked by re-probing the same membrane with anti-CD95L antibody (Abnova, Taiwan). For DNA conjugation, potential disulfide bridges after His-tag cleavage were reduced by addition of TCEP to a final concentration of 2 mM and incubation for 30 min at 4 °C. Afterwards DNA-maleimide (biomers.net GmbH) was added in 5x excess over the CD95L-IZ trimer and incubated overnight at 4 °C. The CD95L-IZ-ssDNA construct was then purified with Amicon centrifugal filters (Merck Millipore, 30 kDa cutoff, cat.no.: UFC5030), with ten PBS washing steps at 8000 rcf and 4°C for 4 min each.

### Native PAGE

To see the band shift of CD95L-IZ after CD95 binding under native conditions, we used clear native PAGE (CN-PAGE) and blue native PAGE (BN-PAGE). Here, we considered CD95L-IZ with a theoretical isoelectric point (pI) of 8.8 as a basic protein and CD95 with a pI of 6.8 as an acidic protein at physiological pH. For CN PAGE, a hand-cast native gel without SDS (7.5 % acrylamide, 0.1 % ammonium persulfate (APS), 0.1 % tetramethylethylenediamine (TEMED), 375 mM Tris, pH 9.4) was freshly prepared. Protein samples of 100 ng CD95-His incubated with 100 ng CD95L-IZ-biotin at 37 °C for 1 h, or 100 ng CD95-His alone (Sino Biological, Beijing, China) were loaded with 10 % glycerol into 10 µl loading volume. Electrophoresis was performed at 4 °C 200 V for 0.5 h with running buffer (50 mM Tris, 192 mM Glycine, pH 9.14). Afterwards, the proteins from the native gel were transferred to the PVDF membrane and probed with an anti-His-tag antibody (BioLegend). For BN-PAGE, protein samples of 100 ng CD95-His alone or with 10x molar excess of CD95L-IZ or CD95L-Enzo were loaded into a precast gel (Biorad) with 50 % glycerol and 5 % Coomassie blue G-250 with a final volume of 10 µl. The electrophoresis was performed with 50 V, 75 V, 100 V gradient voltage each for 30 min at 4 °C with anode running buffer (25 mM Tris/ 192 mM glycine, pH 8.3) and cathode running buffer (25 mM Tris/ 192 mM glycine/ 0.02 % G-250, pH 8.3). Afterwards, proteins from the native gel were transferred to the PVDF membrane and probed with anti-CD95 antibody (Miltenyi Biotec, clone DX2).

### Droplet assay

8 µl of 0.1 mg/ml biotinylated protein CD95L-IZ-biotin was mixed with 3.1 µl of ATTO594 NHS-ester (stock concentration of 18 µM, ATTO-TEC GmbH, Siegen, Germany) and 11.1 µl of the reaction mix was incubated at 4 °C overnight. The next day, 4.3 µl of the above reaction mix was taken and diluted in Phosphate-buffered saline (PBS) at pH 7.4 at a final protein concentration of 10 µg/ml in 31 µl. A drop of the protein solution was placed in one well of an 8-well chamber (Sarstedt) and incubated at r.t. for 1 h. The droplet was pipetted out and shortly air dried to ensure negligible free protein in the solution. Thereafter, the whole chamber surface was covered with 400 µl of 10 µg/ml BSA (ITW Reagents) and incubated at r.t. for 1 h. The BSA droplet was removed and the chamber was washed by 3 x 800 µl PBST buffer (phosphate-buffered saline, 0.1 % tween-20 at pH 7.4) followed by incubation with 400 µl of 10 µg/ml AF488-streptavidin (AAT Bioquest) at r.t. for 1 h. The chamber was washed with 5 x 800 µl PBST afterwards. The sample was imaged with an IX83 inverted microscope (Olympus Europa SE & CO. KG, Hamburg, Germany) using a 20x air objective (NA 0.85, UPLSAP 20x O, Olympus). By fluorescence imaging using suitable filters in the 594 nm and 488 nm excitation channels, colocalization of the biotinylated protein and streptavidin was checked to verify successful protein biotinylation.

### ELISA of CD95 and CD95L-IZ-biotin binding

CD95-His receptor protein (Sino Biological) was coated on a 96 well plate (Sarstedt) at 2 µg/ml concentration in 50 µl PBS buffer overnight at 4 °C. Each well was washed 3 times with 180 µl PBS-0.05 % tween-20 after each incubation step. The next day, wells were washed 3 times and blocked with 180 µl 2 % BSA in PBS −0.05 % tween-20 for 1 h at r.t. CD95L-IZ-biotin or soluble CD95L (Enzo Life Sciences Inc., Lörrach, Germany) was diluted from 0.1 nM to 100 nM in 0.5 % BSA/PBST and incubated with CD95-His at r.t. for 2 h followed by washing. Primary anti-CD95L antibody (Abnova) was diluted into 10 µg/ml in 50 µl 0.5 % BSA/PBST and incubated overnight at 4 °C. The next day, wells were incubated with goat anti-mouse HRP linked secondary antibody (Cell Signaling) with 1:1000 dilution in PBST for 1 h at r.t., each well received 50 µl. Afterwards, 100 µl 1-StepTM slow TMB ELISA substrate (Thermo Fisher Scientific) was added into each well for 30 min for color development. Absorbance was measured at 652 nm with a Tecan Infinite 200PRO (Tecan Life Sciences). Absorbance values were corrected for the background signal and normalized by dividing with the maximum value. Data points were fitted using Matlab (R2018a, The MathWorks, Inc., CA, USA) with a 4-parameter logistic model.

### Apoptosis dynamics assays

Apoptosis dynamics assays were performed to demonstrate the killing efficiency of CD95L-IZ-X (CD95L-IZ-His/CD95L-IZ/CD95L-IZ-biotin) and its inhibition by CD95-His (Sino Biological) or enhancement by anti-His-tag antibody (BioLegend). Soluble CD95L (Enzo Life Sciences Inc.) was assayed for a comparison of the apoptosis induction rate with CD95L-IZ-X. Hela WT cells were seeded in the wells of an 8-well glass slide with thickness No.1.5 (Sarstedt) at least 24 h before the experiment and the DMEM complete culture medium was replaced with pre-warmed 200 µl Leibovitz‘s L-15 medium without phenol red (Gibco) supplemented with 10 % FBS (Gibco) + 1 % P/S (Sigma-Aldrich) just before CD95L-IZ-X addition. CD95L-IZ-X (i) alone or (ii) incubated with CD95-His receptor or with (iii) anti-His-tag antibody (at r.t. for 30 min) were added to the 8-well plate with a final volume of 300 μl in each well at the desired concentration. As a general control, cells on one well were incubated with L-15 complete medium without ligand, where no apoptosis of cells was observed. As another control to exclude unspecific killing, the Hela CD95^KO^ cell line was incubated with the secreted ligand in L-15 medium after 3 days of the expression. Again, no apoptosis was observed. The 8-well chamber was mounted on an IX83 inverted microscope (Olympus Europa SE & CO. KG, Hamburg, Germany) using a 20x air objective (NA 0.85, UPLSAP 20x Olympus) on a temperature-controlled stage heating system (PeCon GmbH, Ulm, Germany) at 37 °C. For each well, phase contrast or bright filed images were taken at 5 positions at 5-15 minutes time intervals over 15 h with an autofocus system using the CellSens Dimensions Software (Olympus). The time-lapse videos were analyzed using Fiji (version 1.49v, U. S. National Institutes of Health, Bethesda, MD, USA). The death time of every single cell undergoing apoptosis, showing membrane blebbing and cell fragmentation, was identified manually. Frame numbers of each apoptosis event were marked individually and exported to a table. This table was loaded with Matlab (R2018a, The MathWorks, Inc.) and fitted with the Hill equation (see equation 1). Here, P_max_ and P_min_ are the maximal and minimal fractions of apoptotic cells, and t_half_ is the time when half of all cells die. n is the Hill coefficient indicating the slope of the curve, corresponding to the speed of the apoptosis signaling. Standard deviations were calculated from measurement replicates.

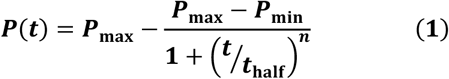

### DNA origami folding & imaging

DNA origami was folded with 12.5 nM p7249 scaffold (tilibit nanosystems), with 4x excess of staples, 8x excess of staples with anti-handles (purchased from IDT) in TAE (40 mM Tris, 20 mM acetic acid, 1 mM EDTA, pH 8.2), supplemented with 12.5 mM MgCl_2_. The folding mixture was heated to 65 °C and subsequently cooled to 4 °C, over 16 h on a Biometra TRIO (analytik jena GmbH+CoKG). The DNA origami was purified with Amicon centrifugal filters (Merck Millipore, 100 kDa cutoff, cat.no.: UFC5100) and washed five times with TAE supplemented with 5 mM MgCl_2_ at 8000 rcf for 4 min each. CD95L-IZ-ssDNA in 5x excess over each binding site on the DNA origami was added to the DNA origami mixture and incubated overnight at 4 °C. The DNA origami conjugated to CD95L-IZ-ssDNA was again purified with Amicon centrifugal filters (Merck Millipore, 100 kDa cutoff, cat.no.: UFC5100) and washed five times with TAE supplemented with 5 mM MgCl_2_ at 8000 rcf for 4 min each.

### Transmission Electron Microscopy

Carbon-coated copper grids (Plano GmbH, cat.no.: S162-3) were glow-discharged with oxygen plasma. Then 10 µl of 2 nM CD95L-IZ-ssDNA on DNA origami were incubated on the grid for 1 min. Excess solution was removed using filter paper. The grid was then stained twice with 10 µl 1.5 % uranyl formate solution. TEM imaging was conducted on a Jeol-JEM-1230 operating at an acceleration voltage of 80 kV. TEM micrographs were analyzed with Fiji version 2.14.0 or later.

## Results

To motivate the CD95L-IZ monomer design, we first introduce the native CD95L structure. **Figure 1A** illustrates the main domains of membrane-bound CD95L monomer. CD95L naturally occurs in a trimeric form, the trimerization being mediated by the THD. A C-terminal domain (amino acids 279-281) is responsible for binding the CD95 receptor. The transmembrane domain enhances the stability of the CD95L trimer, and such stabilization was suggested to play a crucial role in the apoptosis initiation (48). The PRD influences the expression level and stability of CD95L. The stalk region can be cleaved by various metalloproteinases (MMPs) to form soluble CD95L which turned out to be significantly less efficient in apoptosis initiation than the membrane-bound, stabilized trimer of CD95L (14,49). The crystal structure of the CD95L extracellular domain, resolved in previous studies (50), reveals a cone-shaped homotrimer (**Figure 1A right**). Within the trimer, each CD95L monomer adopts a jelly-roll fold with two beta-sheets. Trimerization of CD95L further provides an optimal spatial arrangement of the binding sites necessary for high-affinity interaction with CD95. Up to three CD95 can be recruited to CD95L, each binding to the groove between two CD95L monomers. CD95L further has three N-linked glycosylation sites at positions Asn184, Asn250, and Asn260 (36,51), and two native cysteine residues, Cys202 and Cys233, forming an intracellular disulfide bridge that enhances structure stability. Mutagenesis studies (52) have shown that charged and polar amino acids, such as cysteine (Cys) and asparagine (Asn) predominantly mediate CD95-CD95L interactions. In contrast, interactions between monomer subunits involve van der Waals forces and hydrogen bonds located between the beta-sheets.

The plasmid design is shown in **Figure 1B** The CD95 ligand extracellular domain (amino acids 137-281) is fused with an IZ domain, as the self-trimerization domain to enhance structure stability and ensure efficient apoptosis induction. This is followed by one extra cysteine for site-specific conjugation (such as biotinylation or DNA conjugation). Additionally, we designed CD95L-IZ either for (1) transient transfection or for (2) stable expression in HEK293T cells. In case of (1), transient transfections, CD95L-IZ was equipped with a His-tag and TEV protease sequence for affinity purification and subsequent cleavage of the N-terminal region. A signal peptide was fused before this sequence at the N-terminus, to allow for secretory expression and to increase the yield. As a cost-efficient reagent for transient transfection in comparison with other commercially available reagents, Polyethylenimine (PEI) was used. PEI was previously shown to be suitable for larger-scale transfection with high efficiency (53,54). PEI is a synthetic polymer with high positive charge density in pH-neutral solutions wherefore it forms complexes with DNA for cell intake. For the complete transient transfection protocol see **CD95L-IZ plasmid design for transient transfections** and **Large-scale expression of CD95L-IZ using cell factory** in the **Methods** section. For comparison, in case (2), a HEK293T cell line stably expressing CD95L-IZ was established, wherefore CD95L-IZ was equipped with a Strep-Tag in combination with a PreScission cleavage site for affinity purification and tag removal. A human VEGF leader sequence was fused before this sequence at the N-terminus, to allow for secretory expression. For the complete protocol see **Formation of stable cell line and protein purification** in the **Methods** section. In the following, we focus on case (1), as here several measures were taken to improve the yield (to 350 µg / 1 L culture), while in case of (2) the yield is already high by the use of a stably expression cell line (of 2 mg / 1 L culture).

Cell morphology was checked before transfection and monitored daily afterwards (**Figure 1C upper panel**). Following the transfection, cells changed from a single, separated shape to a clustered, cloud-like shape, and the culture medium became cloudy. Cell culture medium was collected 72 h post-transfection. Naturally, also the number of floating cells (dead cells) increased over time. However, we checked that these were not due to CD95L-IZ induced apoptosis, also since HEK293T cells lack CD95. **Figure 1C lower panel** shows HEK293T cells transfected by PEI/DNA in a 10-layer cell factory in the cell culture hood (left) and cell culture incubator (right). We chose to use the cell factory due to the following advantages: first, it greatly saves time, space, and material in scaling up cell numbers in comparison to the use of, e.g. T175 flasks, since its surface area of 6360 cm^2^, which is equivalent to 36 T175 flasks. It further leads to a lower contamination risk. Second, usage of the cell factory is based on a regular adherent cell culture routine, which does not need additional laboratory equipment. The CD95L-IZ-His expression and secretion in the medium 72 h post-transfection was verified by western blot. It was verified by both, the anti-CD95L antibody (**Figure 1D**) and the anti-His-tag antibody (**Supplementary Figure 1**) resulting in a signal band at ∼ 35 kDa indicating the monomer CD95L-IZ-His. The slight tilt of the signal lane in **Figure 1D** was due to the high FBS amount in the concentrated medium, causing a high viscosity.

To increase the success rate and purity of the CD95L-IZ-His purification, we employed affinity purification using anti-His-tag antibody coupled agarose beads. We obtained a high yield of 455 *μg* from 1.3 L medium of human CD95L-IZ-His, enabling downstream modifications such as His-tag cleavage, biotinylation, DNA hybridization, or fluorescent labeling. The purity and functionalization then allow for subsequent quantitative biophysical studies and applications in nanotechnologies. Agarose beads were used in a batch procedure and elution was achieved by an alkaline elution method based on a calculated high theoretical isoelectric point (pI) of CD95L-IZ-His (pI 8.8). **Figure 2A lane 1** shows the SDS-PAGE result of CD95 ligand purification with a purity of 86 %, based on calculating the integrated peak intensity over the total intensity signal.

**Figure 2.**
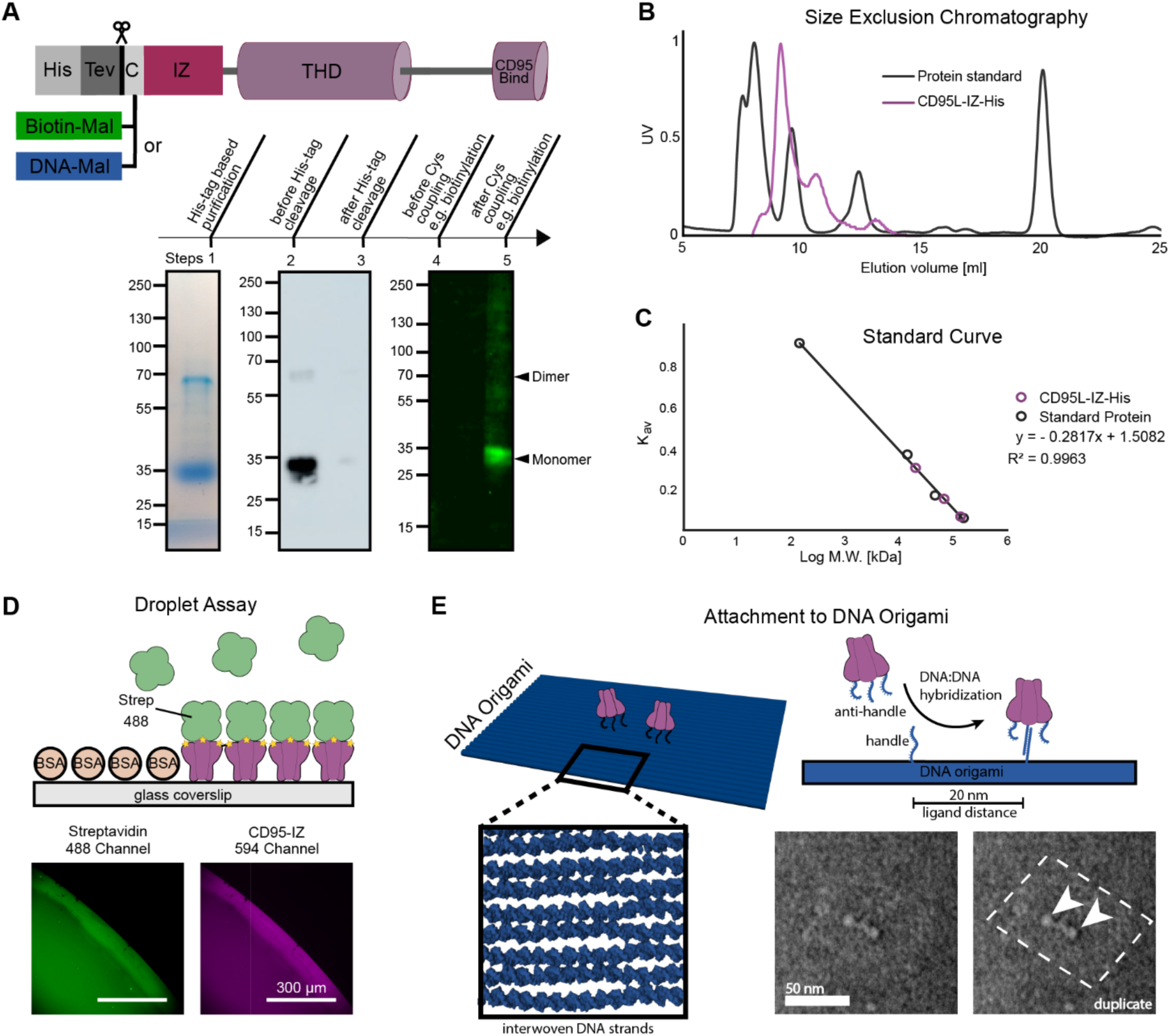
Purification, modification, and characterization of CD95L-IZ. **A** CD95L-IZ purification and functionalization. **Lane 1** Affinity purification of CD95L-IZ-His using anti-His-tag antibody coupled agarose beads followed by 10 % SDS-PAGE analysis of the purified protein. **Lane 2-3** Western blot analysis using anti-His-tag antibody: 100 ng CD95L-IZ-His before (lane 2) and 100 ng CD95L-IZ after His-tag cleavage (lane 3). **Lane 4-5** Western blot analysis of site-specific biotinylation using the cysteine maleimide reaction: 300 ng CD95L-IZ before (lane 4) and after (lane 5) biotinylation were loaded, probed with AF488-streptavidin. **B** Size exclusion chromatography (SEC) analysis of CD95L-IZ-His after His-tag affinity purification compared with a protein standard mix run under identical conditions. The SEC data exhibits peaks corresponding to the trimeric, the dimeric, and the monomeric form of CD95L-IZ-His. **C** From the SEC data, a standard curve of the protein standard mix is calculated and peak positions of the CD95L-IZ-His data are plotted to derive the corresponding molecular weight. **D** Droplet assay to verify biotinylation. Top: a sketch of the assay. Bottom: Fluorescence colocalization images of the 488 nm and the 594 nm channel. CD95L-IZ-biotin was non-specifically labeled with ATTO594 NHS-ester and incubated with AF488-streptavidin on a BSA passivated glass coverslip. **E** DNA origami functionalized with CD95L-IZ-ssDNA. The DNA handle on the CD95L-IZ hybridizes to a complementary anti-handle on the DNA origami, attaching the ligand site-specifically to the DNA origami. The TEM micrograph on the lower right is duplicated and origami outlines and proteins are indicated for better interpretability.

The monomeric molecular weight (M.W.) of the secreted CD95L-IZ without modification (i.e. with His-tag but without the cleaved signaling peptide) is calculated as 24.1 kDa. On SDS-PAGE the monomer CD95L-IZ runs at a molecular weight of 35 kDa due to glycosylation (**Figure 2A lane 1**). N-linked glycans can add substantial mass to the protein, typically ranging from 1-3 kDa per glycan, and even higher in more complex forms. The observed 11 kDa difference suggests the presence of three to five N-glycans in CD95L-IZ. Furthermore, glycosylation is often heterogeneous, resulting in different glycan structures on individual proteins, which manifests itself as a smear or multiple bands in the SDS-PAGE (**Figure 2A lane 1**). Under reduced conditions, the monomeric band at 35 kDa was detected, as expected. Additionally, dimers were observed around 70 kDa, which persisted despite heating and addition of the TCEP reducing agent. These dimers may arise from strong hydrophobic interactions, ionic and hydrogen bonds between CD95L monomers that cannot be disrupted by SDS molecules. The dimers can be observed from previous purification results.(29,32,33). Overall, this observation suggests a particularly stable dimeric structure.

After 3 days of secretion expression, the cell pellet was collected and tested by western blot. There was no CD95L-IZ-His identified in the cell lysate (data not shown here), indicating a high secretion rate. The biological activity of CD95L-IZ-His before purification was assessed by incubating the collected medium with Hela WT cells, estimating the ligand concentration to be approximately 2000 ng/ml (**Supplementary Figure 3**). This was calculated from the signal intensity of the band in western blot analysis (**Figure 1A**) and the CD95L-IZ-His protein concentration in the expression medium ranging from 60 to 125 ng/ml. This concentration estimate turned to be a lower limit, since the apoptosis dynamics curve of the unpurified ligand (**Supplementary Figure 3**) showed an apoptosis induction efficiency of 100% at long time scales. The estimated 2000 ng/ml ligand concentration may hence be higher, with the difference arising from protein loss during the preparation steps. Since CD95L-IZ with and without His-tag differs only by a M.W. of 1.8 kDa, no visible size differences were expected and also observed in the SDS-PAGE and western blot used to test for the His-tag cleavage. After His-tag cleavage (**lane 3**) a significant reduction in the signal intensity in comparison to the signal before cleavage (**lane 2**) is observed, yielding a high calculated cleavage efficiency of 98 %.

**Figure 2B** shows the oligomerization analysis of CD95L-IZ-His using size exclusion chromatography (SEC). Protein standards were analyzed under identical conditions and verified by SDS-PAGE. The elution volumes of standard proteins (listed in **Table 1**) were used to construct the standard curve shown in Figure 2C. Oligomer sizes were calculated and are summarized in **Table 2**, yielding an estimated trimer size of 137 kDa, dimer size of 67 kDa, and monomer size of 20 kDa. Oligomerization of CD95L-IZ-His was further examined using western blot under non-reducing conditions (**Supplementary Figure 1 lane 2),** resolving monomers, dimers, and trimers of CD95L-IZ-His with apparent molecular sizes of approximately 36 kDa, 63 kDa, and 110 kDa, respectively. The size discrepancy observed between the CD95L-IZ-His in SEC and SDS-PAGE may arise from the different shielding of charges of larger complexes in SDS-PAGE or, in SEC, from an increased hydrodynamic volume that proteins with highly non-spherical shapes can exhibit. The protein purity following SEC purification increased to 97 % (**Supplementary Figure 1 lanes 4 and 5**).

**Table 1.** The standard curve is calculated from the listed values of the elution volume and molecular weight of the protein standard mix. With V_e_ = elution volume, V_o_ = column void volume (8 ml), V_c_ = geometric column volume (24 ml), K_av_ = (V_e_ - V_0_) / V_c_-V_0_, molecular weight (M.W., kDa).

| Protein standard | Elution Volume ($V_e$, ml) | $K_{av}$ | M.W., kDa | Log M.W. |
| --- | --- | --- | --- | --- |
| γ-globulins | 8.96 | 0.06 | 150.00 | 5.18 |
| Albumin chicken egg | 10.71 | 0.17 | 44.30 | 4.65 |
| Ribonuclease A | 13.86 | 0.37 | 13.70 | 4.14 |
| pABA (para-aminobenzoic acid) | 22.46 | 0.91 | 0.14 | 2.14 |

**Table 2.**
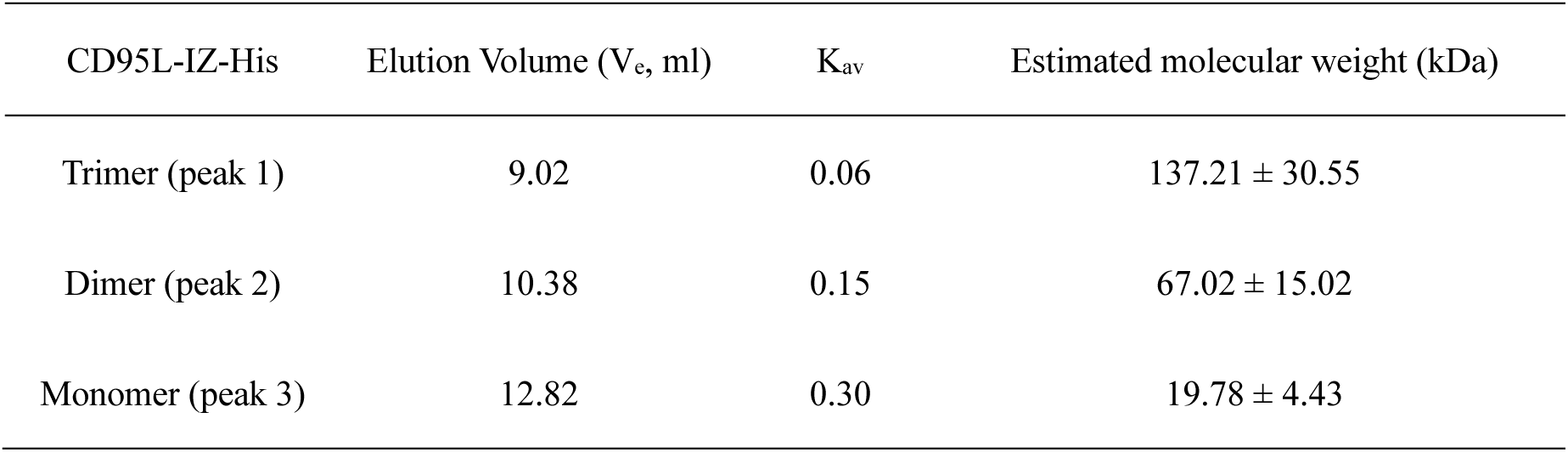
CD95L-IZ-His size estimation from the elution volume following the standard curve of the protein standard mix.

The SEC curve confirms the absence of aggregation in the purified CD95L-IZ-His sample. Additionally, a fit of the SEC curve with three Gaussians (**Supplementary Figure 2**) reveals that CD95L-IZ-His exists as approximately 21 % trimers, 56 % dimers, and 23 % monomers in solution. This distribution is in line with western blot results obtained under non-reducing conditions (**Supplementary Figure 1 lane 2**). While in case of the latter, denaturation effects of SDS might still occur, SEC provides a native oligomerization profile of CD95L-IZ-His. Finally, after cross-linking of CD95L-IZ-His with 4 % formaldehyde (**Supplementary Figure 1 lane 5**), western blot under non-reducing conditions reveals the formation of high-order multimers (indicated by an asterisk) composed of multiple trimers. This phenomenon may be attributed to intermolecular cysteine interaction or a so-called domain-swapping mechanism of isoleucine zipper, as proposed in (55).

CD95L-IZ contains two native cysteines that form intracellular disulfide bridges crucial for its structural stability (52). While the reducing environment of the cytoplasm maintains cysteine residues in their thiol(-SH) form, the additional unpaired cysteine fused at the N-terminus can undergo a non-specific oxidation either during post-secretion or during purification under non-reduced conditions. Therefore, TCEP was used to ensure the accessibility of the thiol group for subsequent maleimide-C5-biotin conjugation. This labeling reaction did not impair the CD95 binding activity or apoptosis induction, as confirmed by subsequent assays (**Figure 3A and 3C**) and is consistent with previous studies (56–58). Biotinylation was initially validated by western blot using AF488-streptavidin, showing the absence of a band for CD95L-IZ without biotin (**Figure 2A lane 4**) and presence of a band after biotinylation (**lane 5**). Biotinylation was further tested by a droplet assay (**Figure 2D**). Here, colocalization of AF488-streptavidin (green) and CD95L-IZ-biotin-ATTO594 (violet) verified the functional and accessible biotin fused to CD95L-IZ.

**Figure 3.**
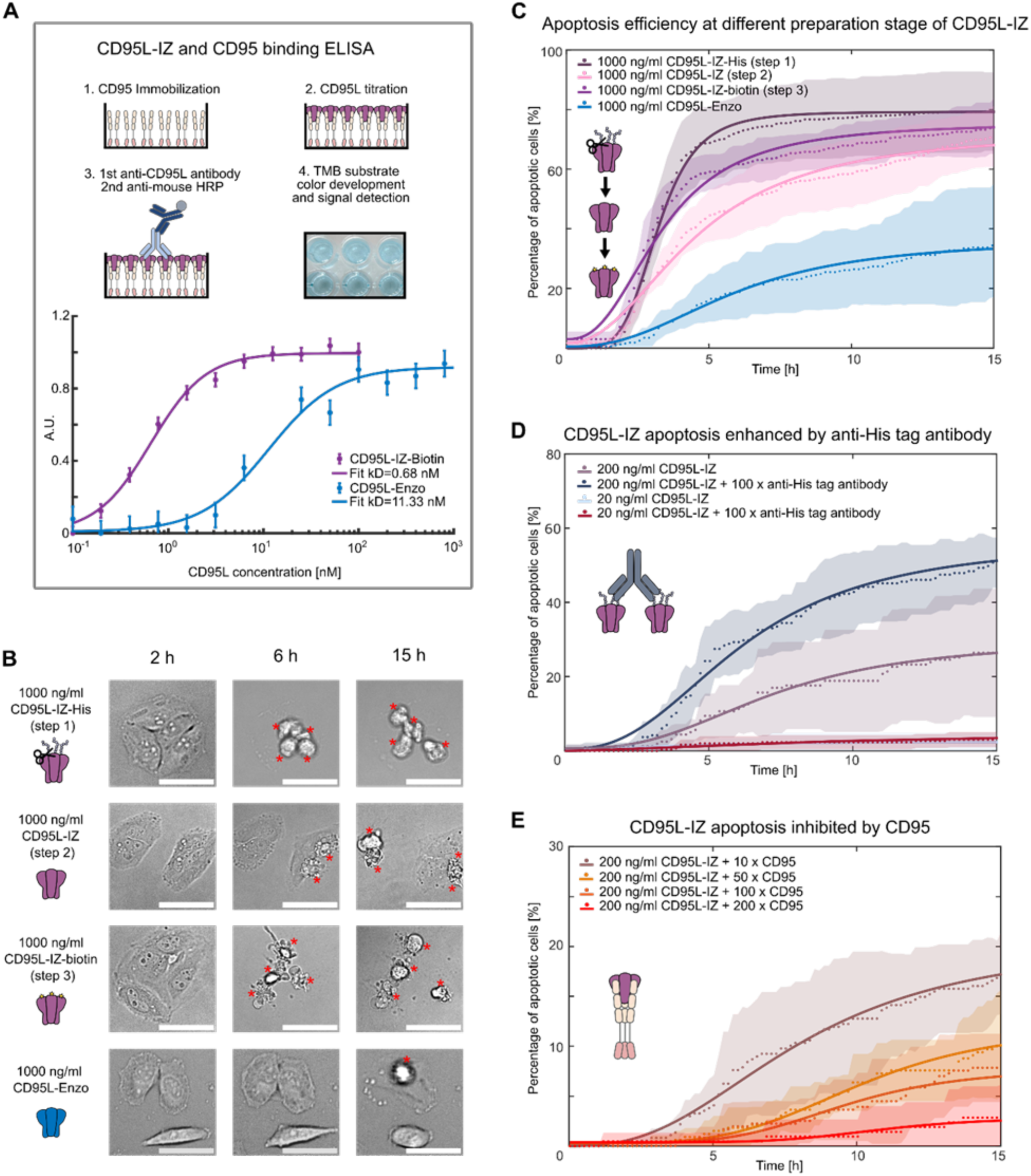
Biological activity assay. **A** ELISA analysis of CD95L-IZ-biotin and CD95L-Enzo binding to CD95. **B** Comparison of representative bright field images showing morphological changes of cells treated with CD95L-IZ-X or soluble CD95L (Enzo Life Science) after 2, 6, and 15 h. Red asterisks indicate the position of apoptotic cell. Scale bars are 50 µm. **C** Apoptosis dynamics analysis of CD95L-IZ apoptosis induction at different stages: post-purification, post-His-tag cleavage, post-biotinylation, and comparison with CD95L-Enzo. **D** The efficiency of CD95L-IZ-His apoptosis induction is increased by anti-His-tag antibody crosslinking. **E** The presence of CD95-His (in 10x, 50x, 100x, and 200x molar excess in comparison to CD95L-IZ-biotin) as competitive binders to the natural CD95 on the cell membrane, successively inhibits CD95L-IZ-biotin apoptosis induction. In panels **C, D, and E** data points are marked with dots. Hill equation fit curves are represented by solid lines, and standard deviations are indicated with shaded regions.

Next to biotin functionalization, we demonstrate the specific attachment of ssDNA to the CD95L-IZ, as confirmed by denaturing SDS-PAGE (**Supplementary Figure 4**) and by attachment to a DNA origami (**Figure 2E**) (59). A ratiometric analysis of band intensities shows, that approximately 2/5 of the CD95L-IZ monomers exhibit a ssDNA modification.

The amount of homotrimers labeled at least once, *P_labeled_*, can then be calculated via the binomial distribution as:

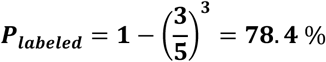

This is the amount of CD95L-IZ-ssDNA homotrimers, which is capable to attach to the DNA origami. The CD95L-IZ-ssDNA homotrimer was then attached to a DNA origami, as depicted in **Figure 2E**. Complementary to the ssDNA sequence on the CD95L-IZ-ssDNA, two ssDNA strands were positioned on the DNA origami with a distance of 20 nm. The CD95L-IZ-DNA origami construct was then verified by TEM imaging (**Figure 2E** and **Supplementary Figure 5**). The probability of CD95L-IZ-ssDNA attachment to the DNA origami was determined by evaluating the TEM micrographs. To avoid inaccuracies that arise with inadvertently discarding badly visible origami, we determined the probability of attachment *p_attached_* by means of stochasticity: the number of DNA origami with one (*N_one_* = 18) and two (*N_two_* = 23) CD95L-IZ-ssDNA attached were counted (**Supplementary Figure 5**) and the respective combinatorial equations

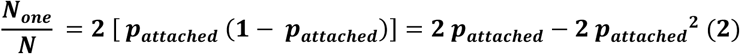

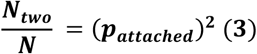

solved for p_attached_, with *N* = *N_one_ + N_two_* = 41. The attachment probability was determined to be ***p_attached_* ≅ 71.2 %.** N.B.: The calculated probability of attachment might change if the DNA origami without proteins could be reliably identified. Further, if the probability of attachment was corrected by the underlying incorporation rate of the respective ssDNA handle into the origami, which is approximately 85 % (59), the probability of attachment for the CD95-IZ-ssDNA would increase to over 80 %.

To verify the biological activity of CD95L-IZ-biotin and soluble CD95L and its binding to the receptor CD95, we first performed enzyme-linked immunosorbent assays (ELISA) or native PAGE analysis, as shown in **Figure 3A** and **Supplementary Figure 6**. From the ELISA binding assay, the dissociation constant (kD) can be derived, with 0.68 nM of CD95L-IZ-biotin and 11.33 nM of CD95L-ENZO. Thus, a more than 10-fold higher binding affinity of CD95L-IZ-biotin in comparison to CD95L-ENZO binding to CD95 is obtained, when applied under identical conditions. This proved that isoleucine zipper indeed increases the trimeric protein stability of soluble CD95L and improves the binding to CD95. In **Supplementary Figure 6A**, the binding of CD95-His with CD95L-IZ-biotin is demonstrated by a band shift in **lane 1,** indicated by the red asterisk. In **Supplementary Figure 6B lane 1,** CD95-His shows the expected monomer signal band at an apparent molecular weight of 44 kDa as well as a broad signal band around 240 kDa, indicating a multimeric form of CD95. This multimeric form occurs at high concentrations under *in vitro* native conditions, e.g. a SDS-stable CD95 complex around 200 kDa was also reported in reference (60). **Lane 2 and lane 3** show the effect of binding CD95L-Enzo or CD95L to CD95-His, which results in slight shifts of the signal bands to higher molecular weights indicated by asterisks.

In **Figure 3C-E**, we employed apoptosis dynamics as a multi-parameter analysis to assess the induction of apoptosis by CD95L-IZ under various stages and conditions. This analysis is instrumental in screening potential anti-cancer drugs or other therapeutic agents as it provides key insights into the onset and rate of apoptosis signaling as well as the final percentage of apoptotic cells, i.e. the efficiency of apoptosis induction. In **Figure 3C** the potency of CD95L-IZ-X to trigger the cell apoptosis pathway was evaluated at different preparation stages: freshly purified CD95L-IZ with His-tag, after His-tag cleavage, after biotinylation, and in comparison to the soluble CD95L from Enzo. Nonspecific cell killing was excluded by control experiments. For example, L-15 complete medium without ligand secretion from HEK293T cells after three days of culture was collected and incubated with Hela WT cells, to rule out the killing effect from other toxic substances expressed by HEK293T cells. As expected, no cell death was observed. Additionally, the secreted ligand in L-15 medium after three days was incubated on Hela CD95^KO^ cell line as a control to exclude non-specific cell killing by other membrane receptor proteins, and no apoptosis was observed in this experiment.

All parameters, obtained from the apoptosis dynamics analysis are shown in **Table 3**. In general, CD95L-IZ-X demonstrated around twice the efficiency in initiating apoptosis compared to soluble CD95L (**Figure 3C** and **Table 3**) at the same concentration. This can be explained by the higher binding affinity of CD95L-IZ with CD95 (**Figure 3A**). As shown in **Figure 3C**, a slight decrease in the apoptosis rate n was observed after His-tag cleavage, from n = 4.7 to n = 2.2. Following biotinylation, the apoptosis rate increased slightly to n = 2.5 (**Table 3**). Similarly, the maximum percentage of apoptotic cells decreased from 79% to 73% after His-tag cleavage and rose again to 76% after biotinylation and the apoptosis half-time increased from 3.2 h to 4.5 h and decreased again to 3.3h (**Table 3**). This data shows, that the integrity of CD95L remains intact during all preparation steps. Yet, CD95L-IZ without the His-tag exhibits a slight decrease in the apoptosis induction efficiency, suggesting a slight destabilization of the CD95L trimer or decrease in the binding affinity to the receptor. Biotinylation, on the other hand, was previously shown to enhance the overall binding efficacy and stability against heat and pH of ligands to their targets (61). We were then interested to see, if further crosslinking of CD95L-IZ-His by anti-His-tag antibody would increase the maximum apoptosis percentage. As shown in **Figure 3D**, while a low concentration of CD95L-IZ-His (20 ng/ml) did not show enhanced apoptosis by antibody crosslinking, a roughly twofold increase in the maximum apoptosis percentage from 29.8 % to 56.7 % was observed at 200 ng/ml CD95L-IZ-His after crosslinking (**Table 3**). This interesting enhancement could be due to several factors: (i) antibody crosslinking may stabilize the ligand and promote a favorable conformation or orientation towards the CD95 receptors. It also (ii) increases the local concentration of CD95L-IZ. Similar effects were previously shown to contribute to a more efficient signal initiation (62,63). Finally, we analyzed the apoptosis initiation of CD95L-IZ inhibited by CD95, when CD95L-IZ was pre-incubated with CD95 at excess concentrations (10x, 50x, 100x, 200x) before measuring apoptosis. Here, the maximum percentage of apoptotic cells was successively reduced from 21 % to 3 % at a 10x and 200x excess of CD95, respectively (see **Figure 3E** and **Table 3**). The reduced efficacy is caused by the sequestration of CD95L-IZ-biotin by excess CD95 and verifies, that CD95L-IZ couples successively more CD95. Here, the added CD95-His acts as a decoy receptor and together with CD95L-IZ-biotin forms inactive complexes that cannot trigger apoptosis anymore.

**Table 3.**
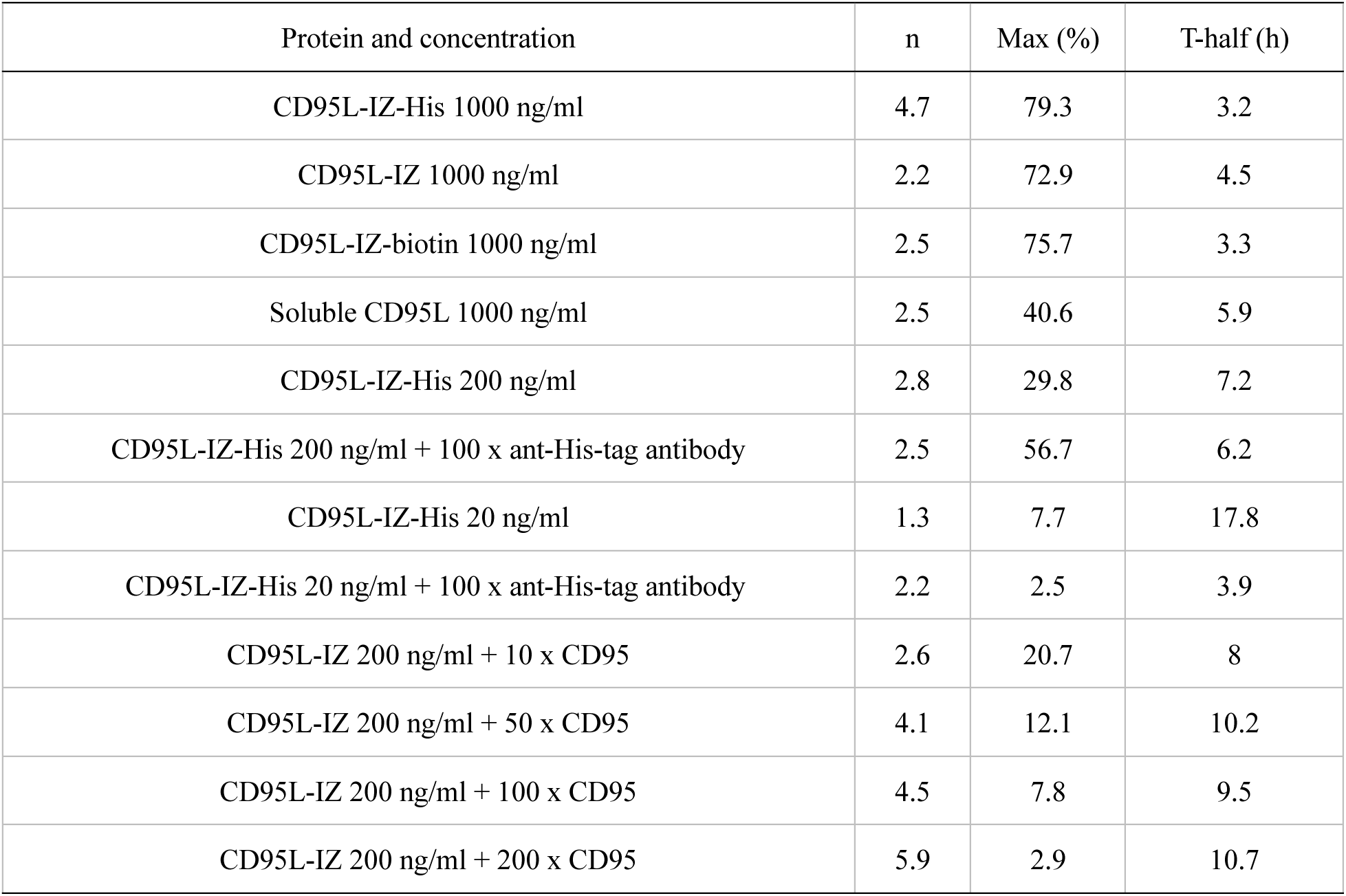
Apoptosis rate (n), percentage of maximum apoptotic cells (Max), and half lifetime (T-half) were obtained from fitting with the Hill equation.

| Protein and concentration | n | Max (%) | T-half (h) |
| --- | --- | --- | --- |
| CD95L-IZ-His 1000 ng/ml | 4.7 | 79.3 | 3.2 |
| CD95L-IZ 1000 ng/ml | 2.2 | 72.9 | 4.5 |
| CD95L-IZ-biotin 1000 ng/ml | 2.5 | 75.7 | 3.3 |
| Soluble CD95L 1000 ng/ml | 2.5 | 40.6 | 5.9 |
| CD95L-IZ-His 200 ng/ml | 2.8 | 29.8 | 7.2 |
| CD95L-IZ-His 200 ng/ml + 100 x ant-His-tag antibody | 2.5 | 56.7 | 6.2 |
| CD95L-IZ-His 20 ng/ml | 1.3 | 7.7 | 17.8 |
| CD95L-IZ-His 20 ng/ml + 100 x ant-His-tag antibody | 2.2 | 2.5 | 3.9 |
| CD95L-IZ 200 ng/ml + 10 x CD95 | 2.6 | 20.7 | 8 |
| CD95L-IZ 200 ng/ml + 50 x CD95 | 4.1 | 12.1 | 10.2 |
| CD95L-IZ 200 ng/ml + 100 x CD95 | 4.5 | 7.8 | 9.5 |
| CD95L-IZ 200 ng/ml + 200 x CD95 | 5.9 | 2.9 | 10.7 |

## Discussion

Soluble CD95L is weakly apoptotic and appears to be outcompeted by membrane-bound CD95L to induce cell death due to its lower stability and lack of oligomerization. For example, an early study suggested, that two CD95L trimers need to form a dimer to induce apoptosis signaling efficiently (19). Other studies reported that biologically functional sCD95L through structural modifications or by adding additional crosslinkers could enhance the apoptosis signal initiation. For example, the fusion of CD95L with a Flag-tag can be crosslinked by anti-Flag-tag antibody (64,65). Other approaches include fusing CD95L with small domains such as tenascin-C (TNC) (20) and T4 fibritin (foldon) (63) to promote self-trimerization. Considering that the geometry and orientation of membrane-bound CD95L may enhance apoptosis signaling, DNA origami has recently been used to create a hexagon geometry for CD95L anchoring (62), significantly enhancing apoptosis efficiency. In addition to stabilizing the trimeric CD95L configuration, a local concentration enhancement of CD95L (e.g. due to coupling on DNA origami platforms) was shown to promote a more efficient signal initiation (62,63). Antibody crosslinking of the CD95 receptor directly was further suggested to maintain the active signaling state(66). Intriguingly, all these studies suggest that spatial fixation of the N-terminus or high-order oligomerization is necessary to amplify the apoptosis signal initiation. Our work shows explicitly, that stabilization of the CD95L trimer via the IZ motif leads to a signaling enhancement, which is further amplified (to a twofold increase in the maximum apoptosis percentage) when CD95L is crosslinked via antibodies (**Figure 3C-D**).

We infer that the mentioned approaches result in a local CD95 concentration enhancement, which in turn recruits larger amounts of adaptor molecules and procaspases 8, facilitating the autocatalytic activation of the latter (67). This in turn could result in large amounts of active caspase 8, a direct activation of effector caspases, and a more direct mitochondria-independent apoptosis pathways, as previously suggested (68,69).

Trimerization of soluble CD95L has been previously confirmed by gel filtration (12) and cross-linking (13), but quantification of the ratio of each oligomer is missing. In **Figure 2C**, we showed the size exclusion chromatography analysis of purified CD95L-IZ-His with quantification of each oligomerization ratio using Gaussian fitting (see **Supplementary Figure 2)**. Thus, we show that while th e majority of the CD95L-IZ occurs in a trimeric state (i.e. 62%), also dimeric and monomeric fraction exists. Care should be hence taken, to account for the different populations, when interpreting the signal induction efficiency. Moreover, site-specific chemical modifications using maleimide-cysteine reaction provide a powerful tool to label the protein. The maleimide cysteine reaction has been considered the most specific labeling reaction due to the high selectivity of the thiol group (-SH). We thus attached CD95L-IZ successfully to DNA origami via complementary DNA-DNA hybridization. Analogously to the biotin modification of the CD95L-IZ, we conjugated a maleimide functionalized ssDNA strand to the cysteine residue on the CDC95L-IZ. The resulting CD95L-IZ-ssDNA conjugate was then attached to a complementary ssDNA on a rectangular DNA origami sheet, used previously for studies on the effects of different CD95L pattern attached to DNA origami on the apoptosis induction (26). The determined probability of attachment for the CD95L-IZ-ssDNA to the handles was ∼ 70 %. This demonstrates the broad applicability of the CD95L-IZ and paves the way for its use in nanoscale therapeutics.

## Conclusion

In this work, we presented a recombinant CD95L exhibiting an IZ motif at the N-terminus, which stabilized the trimerized CD95L. A fast, cheap and high-yield production of this CD95L-IZ *via* the HEK293T secretory expression system is presented, along with CD95L-IZ affinity purification, and characterization of its biological activity and complex formation with CD95. We demonstrated that this stabilization results in a high apoptosis induction efficiency, when cancer cells were exposed to CD95L-IZ in solution in comparison to the unmodified CD95L. A cysteine amino acid fused behind the IZ further served as a versatile coupling site for bionanotechnological or biomedical CD95L functionalization with biotin or DNA hybridization, which can be exploited in future for new anticancer drug developments.

## Ethics

No ethics approval was required for the use of any of the cell lines in this study. The cell lines used in this study were Hela wild type cells and HEK293T cells, which are available from the ATCC organization (Manassas, VA, USA).

## Availability of data and materials

All data are shown in the main text or the supplementary materials. Plasmids are subject to the Uniform Biological Material Transfer Agreement. Requests for the plasmids should be submitted to Cornelia Monzel.

## Abbreviations

CD95: Cluster of Differentiation 95; TNF: Tumor-necrosis factor; IZ: isoleucine zipper; TNFSF: tumor necrosis factor super family; SFK: src-family kinase; mCD95L: membrane bound CD95 ligand; sCD95L: soluble CD95 ligand; MMP: metalloprotease; TM: transmembrane; THD: TNF homology domain; E.coli: *Escherichia coli;* COS: *Dictyostelium discoideum*; 2 x YT: 2 x yeast extract tryptone; LB: lysogeny broth; SDS-PAGE: Sodium dodecyl-sulfate polyacrylamide gel electrophoresis; HEK293: human embryonic kidney 293; DMEM: Dulbecco’s modified eagle medium; FBS: fetal bovine serum; P/S: penicillin/streptomycin; TBS: tris-buffered saline; TCEP: tris(2-carboxyethyl)phosphine; PBS: phosphate-buffered saline; pABA: p-aminobenzoic acid; TEV: tobacco etch virus; DTT: dithiothreitol; CD95 ECD: CD95 extracellular domain; BS(PEG)5: PEGylated bis(sulfosuccinimidyl)suberate; PVDF: polyvinylidene difluoride; BSA: bovine serum albumin; HRP: horseradish peroxidase; ECL: enhanced chemiluminescence; CN PAGE: clear native polyacrylamide gel electrophoresis; APS: ammonium persulfate; TEMED: tetramethylethylenediamine; TE: tris-EDTA; Sf9/Sf21: *Spodoptera frugiperda* 9/21; CHO: chinese hamster ovary: IL2pept: human interleukin 2; PEI: polyethylenimine; Ni-NTA: nickel-nitrilotriacetic acid; TNC: tenascin-C; Cys: cysteine; Asn: asparagine; Ser: serine; SEC: size exclusion chromatography; FA: formaldehyde; TEM: transmission emission microscopy; pI: isoelectric point; ELISA: enzyme-linked immunosorbent assay; TMB: 3,3’,5,5’ - tetramethylbenzidine; CRISPR: clustered regularly interspaced short palindromic repeats;

## Authors’ contributions

Conceptualization: CM, XS

Methodology: XS, NB, JMW, SS, JB

Validation: CM

Formal analysis: CM, XS

Investigation: CM, XS

Resources: CM, AHJ

Writing - original draft: CM, XS

Writing - Review & Editing: NB, JMW, AHJ

Visualization: XS, JMW

Supervision: CM, AHJ

Project administration: CM

Funding acquisition: CM

## Acknowledgments

We thank Prof. Alexej Kedrov (Synthetic Membranesystems, Heinrich-Heine University) for providing the Amersham Western Blot Imager and the group of Prof. Claus Seidel (Molecular Physical Chemistry, Heinrich-Heine University) for providing laboratory space and lab equipment. We thank Joël Beaudouin (formerly IBS, Grenoble) for kindly providing the Hela CD95^KO^ cell line. We thank the protein core facility and Barbara Steigenberger at the mass spec core facility of the Max Planck Institute of Biochemistry for large scale synthesis of CD95L-IZ and subsequent characterization.

