## Supplementary Information for "High yield purification of an Isoleucine zipper modified CD95 Ligand with either biotin or DNA-oligomer binding domain for efficient Cell Apoptosis Induction"

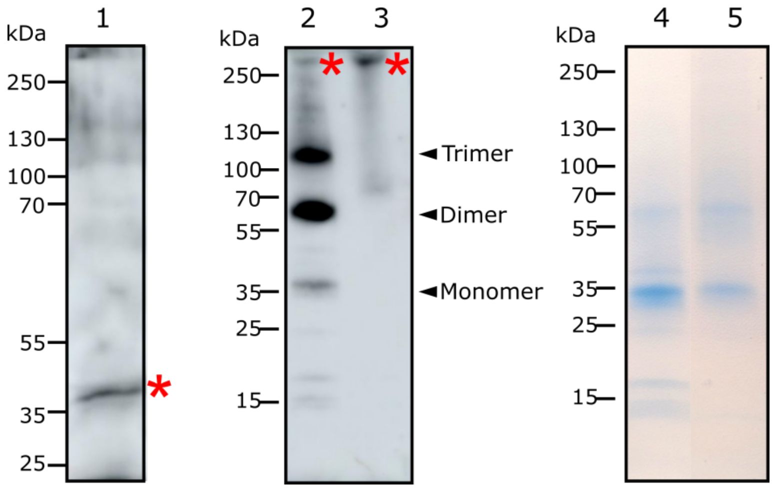


**Supplementary Figure 1. Lane 1:** Detection of CD95L-IZ-His expression by western blot, using an anti-His-tag antibody. **Lane 2:** Western blot analysis of non-reduced CD95L-IZ-His, probed by anti-His-tag antibody. The quantification based on intensity ratios shows a trimer at 35.5 %, a dimer at 50 %, and a monomer at 14.5 % (ImageJ quantification). **Lane 3:** CD95L-IZ-His cross-linked by 4 % formaldehyde (FA), reveals the presence of a high-order oligomer (indicated by asterisks). **Lane 4 and Lane 5:** Purity comparison of CD95L-IZ-His before and after size exclusion chromatography, with observed purities of 86 % and 97 %, respectively.

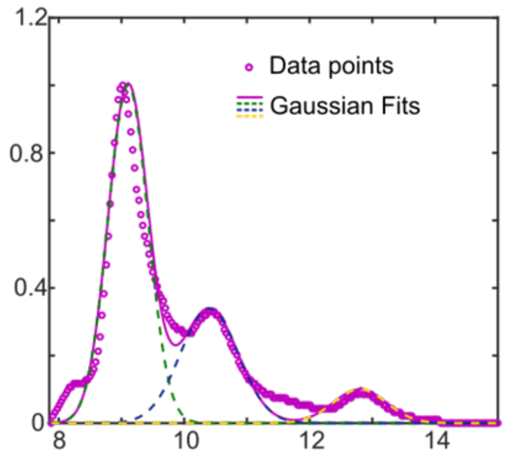


**Supplementary Figure 2.** Size exclusion chromatogram of CD95L-IZ-His, fitted with three Gaussian curves. The trimer is indicated by a dotted green line, the dimer by a dotted blue line, and the monomer by a dotted yellow line. The overall fit is depicted by a purple line, which is the sum of the three Gaussian functions. Data points are shown as purple dots. The fitting results are as follows: Gaussian 1: amplitude = 1.00, mean = 9.10, standard deviation = 0.45, area = 0.80; Gaussian 2: amplitude = 0.34, mean = 10.40, standard deviation = 0.65, area = 0.39; Gaussian 3: amplitude = 0.10, mean = 12.80, standard deviation = 0.60, area = 0.11. The percentage of different oligomers can be calculated as 62 % trimer, 30 % dimer, and 8 % monomer.

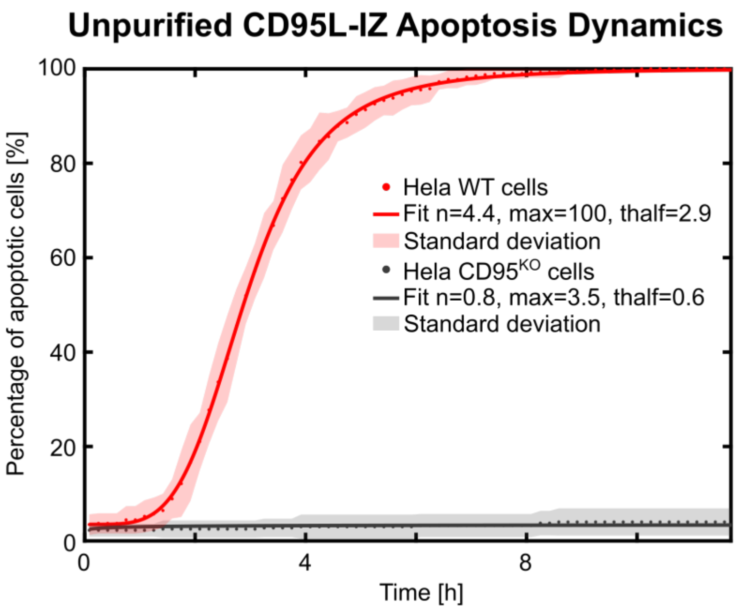


**Supplementary Figure 3.** Functionality assay of CD95L-IZ-His before purification on Hela WT and Hela CD95$^{KO}$ cell lines. A total of 300 µL of L15 complete medium, containing CD95L-IZ-His secreted after 3 days of transfection in HEK293T cells, was incubated overnight with the HeLa WT (shown in red) or Hela CD95$^{KO}$ (shown in gray) cell line. Data points were fitted with the Hill equation with a result for Hela WT: the Hill coefficient (n) = 4.4, the maximum percentage of apoptotic cells (max) = 100 %, and the halflife-time (thalf) = 2.9 h; for Hela CD95$^{KO}$: the Hill coefficient (n) = 0.8, the maximum percentage of apoptotic cells (max) = 3.5 %, and the halflife-time (thalf) = 0.6 h. The graph reaches a value of 100 % apoptotic cells, which typically occurs at high ligand concentrations, here estimated for the unpurified CD95L-IZ-His to 2000 ng/mL. As a negative control, the Hela CD95$^{KO}$ cell line showed negligible apoptosis events.

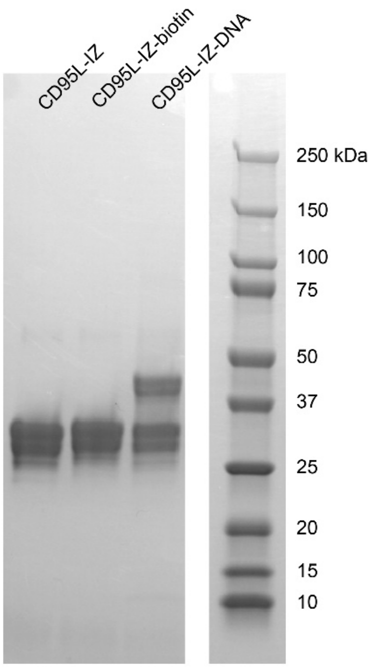


**Supplementary Figure 4.** Denaturing SDS-PAGE gel of CD95L-IZ functionalized with biotin or DNA in comparison with non-functionalized CD95L-IZ.

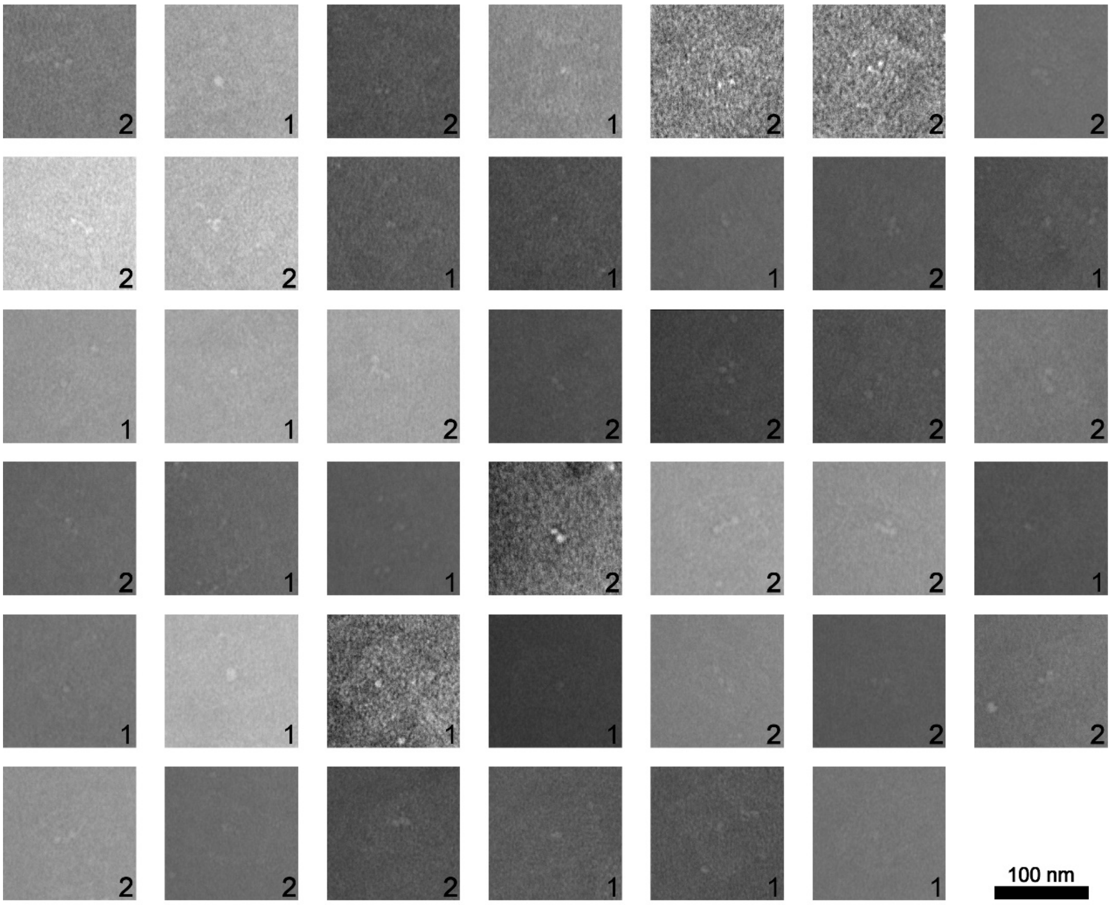


**Supplementary Figure 5.** Montage of cropped TEM micrographs of DNA origami with CD95L attached to them. The number of FasL on the respective DNA origami is written in the lower right corner of every cropped image. For better visibility, the contrast was enhanced in some micrographs. The probability of CD95L attachment is ~ 70 %.

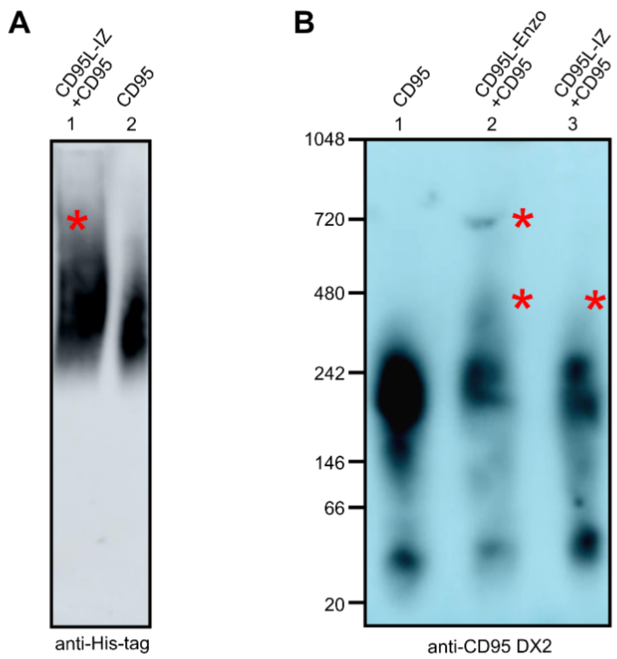


**Supplementary Figure 6. Native PAGE analysis of CD95L-IZ or CD95L-ENZO binding to CD95. A** Clear native PAGE analysis of CD95L-IZ-biotin binding to CD95. Lane 1: 100 ng of CD95L-IZ-Biotin was incubated together with 100 ng CD95 (Sino Biological, His-tagged), and lane 2: 100 ng CD95 alone. The membrane was probed with an anti-His-tag antibody (Biolegend) to detect the shift in the CD95 signal band (indicated by an asterisk symbol). **B** Blue native page analysis of CD95L-IZ-biotin binding to CD95. The blot was probed with anti-CD95 antibody (Miltenyi Biotec, clone DX2), with lane 1: 100 ng CD95 alone (Sino Biological, His-tagged), lane 2: 100 ng CD95 incubated with 10x molar excess of CD95L-Enzo, and lane 3: 100 ng CD95 incubated with 10x molar excess of CD95L-IZ. Bands at higher molecular weight, indicated by asterisks, show fractions of CD95 forming complexes with CD95L-ENZO or CD95L-IZ. Unstained protein standard (Invitrogen) was used as molecular marker.

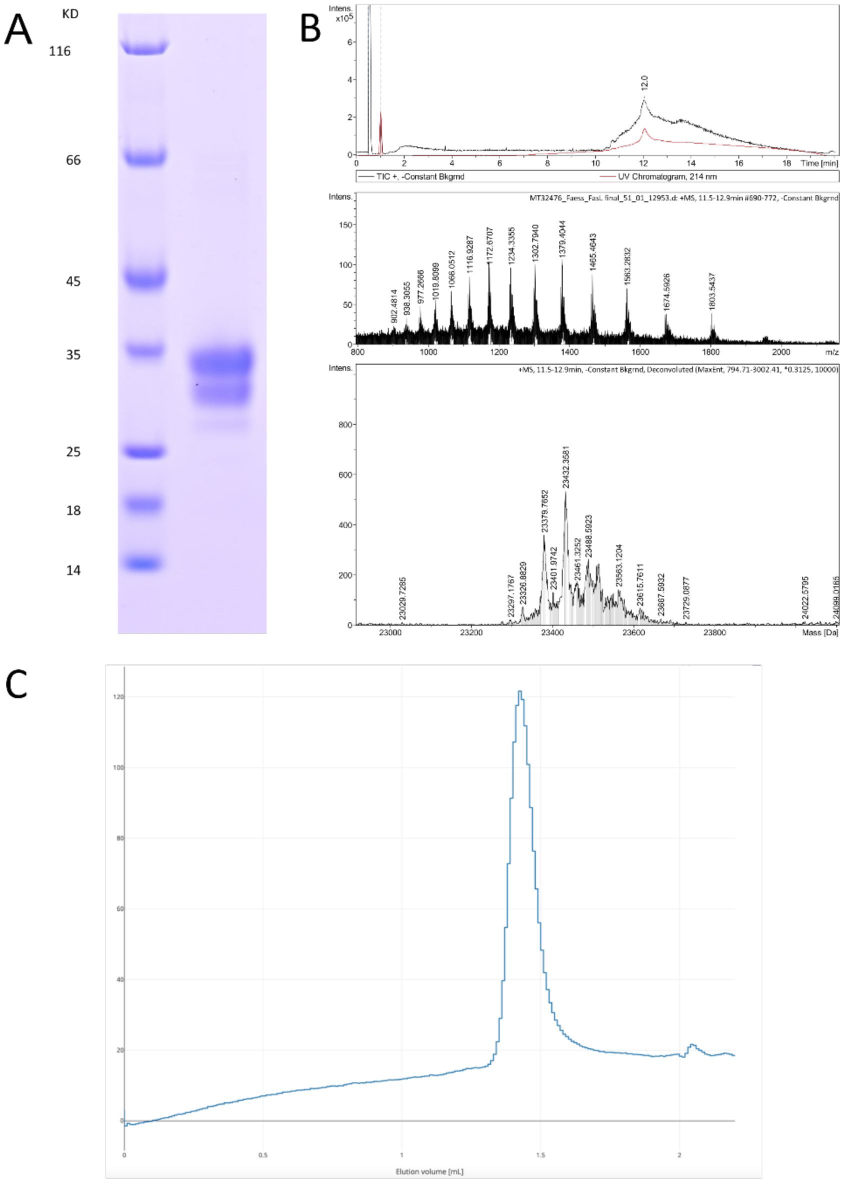


**Supplementary Figure 7.** Quality control of CD95L-IZ produced from the stable cell line (method 2). **A.** SDS-PAGE analysis with gradient gel 4-12 %. Lane 1: molecular weight marker, Lane 2 purified CD95L-IZ. The purified protein runs as a double band reflecting different glycosylation states. **B.** Liquid chromatography-mass spectrometry (LC-MS) analysis with the intact mass of the purified batch. The mass spectrometry spectra confirm different glycosylation states. **C.** Analytical gel filtration with a Superdex™ 200 increase 3.2/300 GL column in a buffer containing 20 mM Hepes (N-2-hydroxyethylpiperazine-N'-2-ethanesulfonic acid) pH 7.5, 150 mM NaCl, and 2 mM DTT.
